# Mechanistic insights into Rho/MRTF inhibition-induced apoptotic events and prevention of drug resistance in melanoma: Implications for the involvement of pirin

**DOI:** 10.1101/2024.09.19.614009

**Authors:** Bardees M. Foda, Annika E. Baker, Łukasz Joachimiak, Marzena Mazur, Richard R. Neubig

## Abstract

**Aim:** Overcoming therapy resistance is critical for effective melanoma control. Upregulation of Rho/MRTF signaling in human and mouse melanomas causes resistance to targeted therapies. Inhibition of this pathway by MRTFi, CCG-257081 resensitized resistant melanomas to BRAF and MEK inhibitors. It also prevented the development of resistance to vemurafenib (Vem). Here, we investigate the role of apoptosis and the protein pirin in CCG-257081-mediated suppression of drug resistance.

**Methods:** Using naïve and resistant mouse YUMMER melanoma cells, we studied the effect of the BRAF inhibitor Vem with or without CCG-257081 on real-time growth and apoptosis (activation of caspase, Propidium iodide (PI) staining, and PARP cleavage). The effects of CCG-257081 on proliferation (Ki67) and caspase-3 activation were assessed in resistant YUMMER_R tumors *in vivo*. Finally, two CCG-257081 enantiomers were tested for pirin binding, inhibition of the Rho/MRTF-mediated activation of ACTA2 gene expression in fibroblasts, and the prevention of Vem resistance development by YUMMER_P cells.

**Results:** Vem reduced growth of parental but not resistant cells, while CCG-257081 inhibited both. The combination was more effective than Vem alone. CCG-257081, but not Vem, induced activation of caspase-3 and -7 in resistant cells and increased PARP cleavage and PI staining. CCG-257081 reduced proliferation and activated caspase-3 in YUMMER_R melanoma tumors. Both CCG-257081 enantiomers robustly suppressed development of Vem-resistant colonies with the S isomer being more potent (1 μM IC_50_).

**Conclusion:** CCG-257081 appears to target pre-resistant cells and Vem-induced resistant cells through enhanced apoptosis. Inhibition of pirin or the Rho/MRTF pathway can be employed to prevent melanoma resistance.

## INTRODUCTION

A critical problem in cancer treatment with targeted therapies is the rapid development of drug resistance. Even immunotherapies are sometimes subject to resistance development^[1]^. Targeted therapies for the aggressive skin cancer, cutaneous melanoma, include drugs that inhibit the MAP kinase pathway, such as BRAF inhibitors (BRAFi), including Vem and dabrafenib, and mitogen-activated protein kinase kinase inhibitors (MEKi), such as trametinib^[2,3]^. Since almost half of melanoma tumors have BRAF^V600^ mutations, the use of these MAPK pathway inhibitors has greatly improved clinical responses, however, drug resistance almost inevitably develops within months. This may be intrinsic resistance in clinical non-responders or induced resistance which develops after an initial response^[4–6]^. So, mechanisms to prevent or reverse resistance would be highly beneficial.

Some resistant melanoma tumors have re-activating mutations in MAPK pathway genes (NRAS, BRAF) and others have mutations in PI3K or RAC1^[7]^. We found that approximately half of the resistant human cutaneous melanomas have increased activation of the Rho/MRTF and/or YAP/TAZ pathway and that these mechanisms appear to play an important role in resistance to targeted therapy in this subset of melanomas^[8,9]^. These pathways are master regulators shaping cytoskeleton architecture and control various cellular processes, including differentiation, migration, proliferation, and focal adhesion^[10,11]^. We previously showed that a series of compounds that inhibit the Rho/MRTF pathway, including CCG-257081, could restore BRAFi-sensitivity in human and mouse melanoma cells^[8,12]^. Moreover, CCG-257081 was also able to prevent the development of Vem-resistant melanoma cells *in vitro*^[9]^.

In addition to its actions to block the MRTF/SRF pathway, CCG-257081 and analogs bind to the protein pirin^[13]^. Pirin is an NRF2-induced protein that has been implicated in melanoma^[13–19]^ While the mechanisms of pirin actions are poorly understood, pirin binding by the MRTF-inhibiting CCG compounds, may be relevant to their anti-melanoma effects. Indeed, pirin has recently been implicated as an inhibitor of apoptosis in colorectal cancer cells^[20]^.

Here, we investigate potential mechanisms of the pirin-binding, MRTF pathway inhibitor CCG-257081 and its actions on drug resistance in melanoma. For these studies, we used the YUMM mouse melanoma cell lines that are isogenic with C57Bl/6 mice that we previously used to assess immunotherapy potentiation by the compound. CCG-257081 inhibits the growth of both Vem-sensitive and Vem-resistant melanoma cells. We also show here that CCG-257081 induces apoptosis. In Vem-sensitive melanoma cells, the combination of Vem and CCG-257081 produces robust and sustained apoptosis. This may contribute to its ability to prevent the *in vitro* development of resistance to Vem. Furthermore, i*n vivo* CCG-257081 treatment of mice carrying Vem-resistant tumors resulted in an increase in caspase activation and attenuated cell proliferation, which likely contributed to the reduced tumor growth reported previously^[21]^. Finally, the two stereoisomers of racemic CCG-257081 showed different activities. The S isomer has a higher affinity for pirin binding, is more potent in inhibiting Rho-dependent transcriptional signaling by the profibrotic cytokine TGFβ, and is more potent at suppressing the development of Vem resistance *in vitro*. These data suggest a role for these compounds and potentially their actions through pirin as a novel combination therapy with MAPK-pathway inhibitors for the initial treatment of melanoma.

## METHODS

### Cell culture

The YUMMER1.7D4 (RRID: CVCL_A2BD) mouse melanoma cell line was purchased from Millipore Sigma (YUMMER1.7D4: cat no. SCC243). Cells were maintained and all experiments performed in DMEM-F12 medium (ATCC no.30-2006) supplemented with 10% fetal bovine serum (Gibco no.10437-028), 1% NEAA (Gibco no.11140-50), and 1% Pen-Strep (ThermoFisher no.15140122). YUMMER1.7D4 is abbreviated as YUMMER_P, where P stands for parental, referring to Vem-sensitive cells. The Vem-resistant YUMMER line, YUMMER_R (R for resistant), was generated as described previously^[12]^. All experiments were performed with mycoplasma-free cells.

### Compounds and Antibodies

Vemurafenib (AmBeed Inc. no. A116840) and CCG-257081 (racemate synthesized in the MSU Medicinal Chemistry Core) were stored as 10 mM stocks in DMSO. Antibodies for PARP (cat no. 9542) and GAPDH (cat no. 5174) were purchased from Cell Signaling. CCG-257081 enantiomers were prepared as described below. The pirin fluorescence polarization reporter^[22]^ was synthesized by the Vahlteich Medicinal Chemistry core at the University of Michigan. Donkey anti-Rabbit800 (cat no. C926-32213) and Donkey anti-Rabbit680 (cat no. 926-68073) immunoblotting secondary antibodies were purchased from LI-COR.

### Incucyte Live-Cell Imaging

The IncuCyte S3 platform (Sartorius) was used to monitor in real-time the activation of caspase-3/7 and cell apoptosis. YUMMER_P and YUMMER_R cells were seeded in 96-well plates (1000 cells per well) and left to adhere overnight. Cells were treated with 5 µM Vem, 10 µM CCG-257081, 5 µM Vem + 10 µM CCG-257081, or 0.15% DMSO (vehicle control in equivalent volume to Vem+CCG-257081). Additionally, cells were concurrently incubated with 1 µM Green caspase-3/7 probe (Sartorius no. 4440) and 0.5 µg/mL propidium iodide (Sigma no. 537060). Untreated cells were unstained and used as a reference for cell growth. Cells were imaged using green, red, and a phase contrast channel. Images were captured every 2 hours for 72 hours using a 10x objective. Captured images were analyzed by IncuCyte software (version: 2021C). The basic analyzer module was applied, and parameters were set to quantify the number of green (active caspase) and red (PI) objects per well. The module also uses an artificial intelligence confluence estimator.

### Apoptosis assessment – PARP cleavage

In addition to the staining for caspase activity and PI staining, to biochemically assess apoptosis, we also assessed PARP cleavage. YUMMER_P or YUMMER_R cells (150,000 cells) were plated into 6-well plates. DMSO or active treatments (10 μM CCG-257081, 5 μM Vem, or both) were applied and cells were harvested 24 hours after the initiation of treatment. Cell lysates were tested by western blot to assess the levels of PARP cleavage.

### Western Blot Analysis

Adherent cells were cultured and treated as indicated. Cells were lysed on ice using lysis buffer (20 mM tris-HCl, pH7.5; 150 mM NaCl; 1 mM NaF; 1 mM ETDA; 0.5% NP40; 0.5 mM DTT) and supplemented with a protease inhibitor (Thermo Scientific no. A32961, Waltham, MA, USA). Ice-cold, whole-cell lysates were sonicated gently with a probe sonicator. An equivalent amount (∼30 µg) of each cell lysate was boiled in an SDS-loading buffer for 10 min. Samples were loaded onto a 10% polyacrylamide gel and transferred to the Immobilon-FL PVDF Membrane (Millipore Sigma, no. IPFL00010). Membranes were blocked in Intercept LI-COR blocking buffer (PBS: 927-70001) and then incubated with primary antibody overnight at 4 °C. Washed membranes were incubated with the appropriate secondary antibody for 1 hour at RT. The immunoblot membrane was washed, dried, and imaged on an LI-COR Odyssey FC imaging system.

### Immunohistochemistry

We performed immunohistochemistry studies on tissues collected from animal experiments that we performed recently^[23]^. Melanoma tissues were collected from YUMMER_R tumor-bearing mice; the experiment is described in Foda, B.M. et al. 2024^[23]^. Briefly, tissues were collected from mice treated for 12 days with a daily intraperitoneal injection of 100 mg/kg CCG-257081 or an equivalent volume of vehicle containing 5% DMSO and 95% polyethylene glycol-400. Formalin-fixed paraffin-embedded tumors were sectioned into 4 µm sections, and unstained tissue slides were prepared by the histology core at MSU. The unstained slides were incubated at 55°C overnight. Immunohistochemistry staining was performed following the recommended manufacturer’s protocol for paraffin-embedded tissues. Briefly, tissue sections were deparaffinized, rehydrated, and heated in 10 mM citrate buffer of pH 6.0 to unmask antigens. Inactivation of endogenous peroxidase was done by treating the tissues with 3% H_2_O_2_, followed by blocking with 1% BSA and 5% normal goat serum. Tumor sections were incubated overnight with primary antibodies specific for cleaved caspase 3 (Cell Signaling no. 9664) or Ki-67 (Abcam no. ab15580). Tumor sections were washed and incubated with goat anti-rabbit IgG, HRP-linked antibody (Cell Signaling no. 7074). Subsequent staining procedures followed Vector Laboratories’ instructions for the HRP/DAB detection kit (SK-4100). Sections were counterstained with Mayer’s Hematoxylin (Sigma no. MH532), then dehydrated and mounted using Permount mounting media (ThermoFisher no. SP15-100). Images were acquired using Olympus BX41 Fluorescence Microscope. For quantification, a minimum of five fields of 40x captured images per tissue were analyzed blindly using ImageJ software (version: 2.14.0/1.54f).

### Generation of CGG-257081 enantiomers

CCG-257081 was produced by the University of Michigan Vahlteich Medicinal Chemistry Core as previously described^[24]^. The R and S enantiomers of CCG-257081 were obtained by chiral separation of the racemate using a Lux® 5 µm Amylose-2, 250x21.2 mm column, with a 10-90% gradient of two solvents: A: Hexane, B: iPr-OH/MeOH 7/3 (v/v) at a flow rate of 15 ml/min over 30 min. The absolute configuration was determined by comparison of the chiral column analytical data for the separated enantiomers with the product obtained via independent synthesis, starting from commercially available (R)-1-[(tert-Butoxy) carbonyl]-5,5-difluoropiperidine-3-carboxylic acid using the synthetic route described in WO2023122325A2 using the mixed anhydride method in the first coupling stage to avoid racemization. The two enantiomers were checked for optical purity using analytical HPLC with a chiral column (Lux® 5 µm Amylose-2, 150 x 4.6 mm) with a gradient of 1−90% of iPr-OH in hexane and a flow rate of 1 ml/min over 30 min. Retention times were CCG-257081 (S enantiomer): 15.6 min and CCG-257081 (R enantiomer): 26.2 min.

### Pirin protein binding assay

Pirin binding of the CCG-257081 racemate and the stereoisomers was tested by competition with a fluorescence polarization reporter compound as described previously^[22]^.

### *In vitro* activity of pirin enantiomers

The biological activity of the CCG-257081 enantiomers was tested in two assays. First, SRE-Luciferase activity was measured to assess MRTF/SRF-regulated gene transcription. This was the assay originally used to identify CCG-1423, the first lead in this chemical series.^[25]^ Briefly, HEK293T cells were transiently transfected with the Luciferase reported plus constitutively active Gα12 subunit, which activates Rho GTPases (RhoA and RhoC) and downstream MRTF/SRF-mediated gene transcription was measured by Luciferase activity as previously reported^[13,25]^. In the second assay, normal human lung fibroblasts (NHLF, CC-2512, Lonza) were stimulated with TGFβ for 24 hours, and the expression of ACTA2 mRNA was determined by qRT-PCR as reported^[13,26]^. Data are averages of two experiments, each performed in triplicate.

### *In Vitro* Clonogenicity Assay

Exponentially growing YUMMER_P cells were harvested. Two thousand cells per well were plated into 6-well plates and left to adhere overnight. On the next day, drug treatment was applied (5 µM Vem ± 1 µM, 3 µM, or 10 µM of each CCG-25081 enantiomer). Plates were monitored until wells treated with Vem alone showed formation of sufficiently large colonies (about 50 cells per colony), which took about 15 days. Wells were washed with PBS and stained with a fixation-staining solution containing 3.7% formaldehyde and 0.5% crystal violet for 30–60 min. Plates were scanned, and ImageJ software (version: 2.14.0/1.54f) was used to count colonies and measure mean area quantification considering colonies with pixel^2^ size at 50–infinity and circularity of 0.2– _1.0_^[27]^.

### Statistical analysis

Statistical analysis was performed as indicated by unpaired two-tailed t-tests for comparative analysis between two groups. A two-way ANOVA test was used for the tumor size analysis. A log-rank test was used for the analysis of mouse survival. Data are presented as mean ± S.E.M, and a p-value < 0.05 was considered statistically significant. All statistical analyses were performed using the GraphPad Prism version 10.1 software (La Jolla, CA).

## RESULTS

### Differential sensitivity to BRAFi and MRTF Pathway inhibition in resistant vs sensitive YUMMER cells

Real-time analysis of cell confluence was performed in the Incucyte system [Figure 1] to assess the effects of VEM and CCG-257081 on YUMMER cell growth. The results confirm those from a previous study^[12]^, which was obtained at single time points at 72 hours with an ATP-detection readout (CellTiter-Glo). The resistant cells (YUMMER_R) grew more slowly than did the parental (YUMMER_P) cells. The resistant cells, as expected, showed less inhibition of growth, with Vem even at a very high concentration (5 μM). Also, as noted previously for both human and mouse melanoma cells^[12,28]^, the Vem-resistant cells were more sensitive to single-drug treatment with the pirin-binding MRTF-pathway inhibitor (CCG compound) than were the sensitive cells. We previously attributed this to the upregulated Rho/MRTF pathway. For both parental and resistant cells, the combination of Vem+CCG-257081 was more effective than Vem alone.

**Figure 1.**
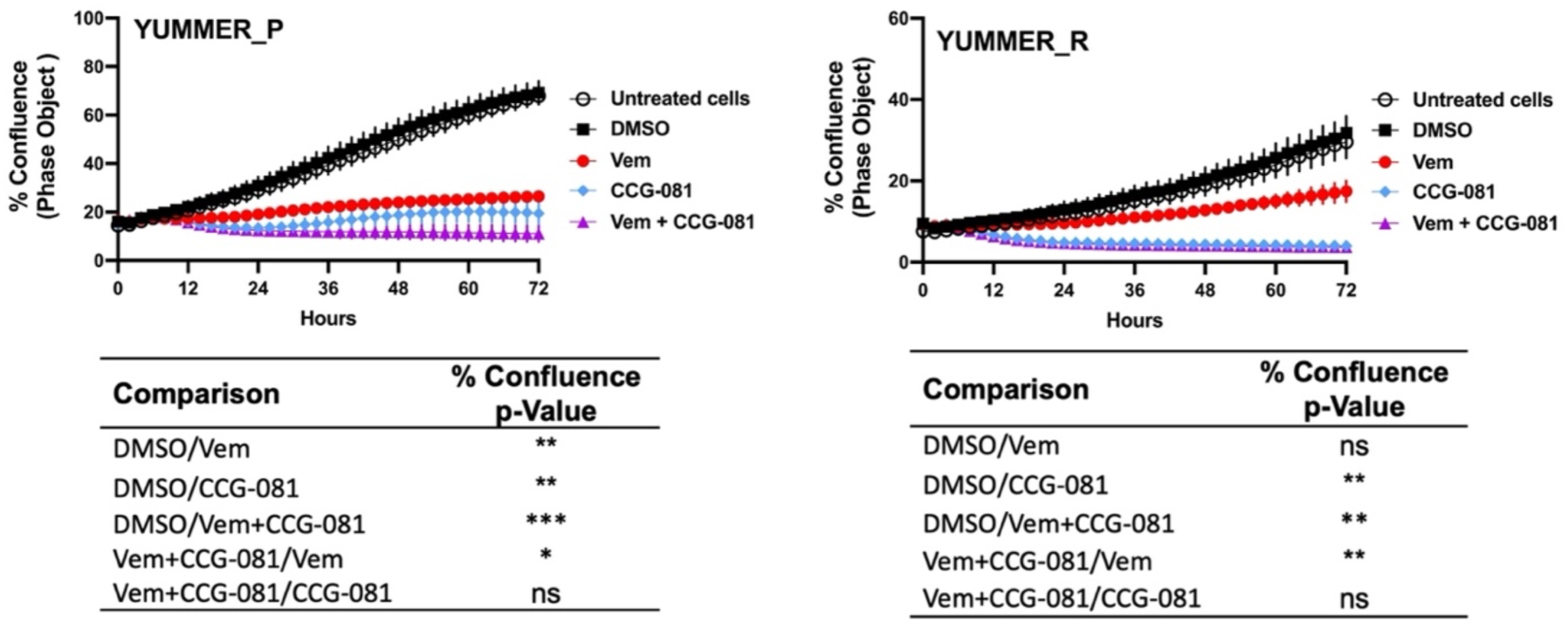
Inhibition of Rho/MRTF pathway enhances Vem-mediated growth suppression of Vem-resistant melanoma cells. Vem-sensitive melanoma cells, YUMMER_P, and Vem-resistant cells, YUMMER_R, were plated at 1000 cells per well in 96-well plates, and growth was monitored in real-time using the Incucyte S3 live-cell imaging system. The indicated compounds (DMSO control, 10 µM CCG-257081 labeled CCG-081, 5 µM Vem, or both 5 µM Vem and 10 µM CCG-081) were added, and the confluence of the YUMMER cells was calculated using the IncuCyte phase object module. Results are the mean ± S.E. of four independent experiments, * p < 0.05; ** p < 0.01; **** p < 0.0001 by A two-way ANOVA test; ns: not significant.

### Combined inhibition of the Rho/MRTF pathway and BRAF produces sustained apoptosis

Both Vem and CCG-257081 prevent the growth of the Vem-sensitive YUMMER_P cells [Figure 1], but the combination treatment causes a major decrease from the initial cell density (as assessed by cell confluence). In contrast, the resistant cells showed only a modest effect of Vem, but CCG-257081, with or without Vem, markedly reduced cell numbers.

To understand a potential mechanism of this effect, we assessed real-time apoptosis in the treated YUMMER_P and YUMMER_R cells. This took advantage of the capabilities of the IncuCyte system and its real-time green-fluorescent caspase-3/7 detection kit with parallel staining of cells with the apoptosis marker, propidium iodide [Figure 2 A and B]. Surprisingly, the parental melanoma cells, YUMMER_P, showed minimal caspase activation with Vem alone, even at a high concentration (5 µM). In contrast, 10 µM CCG-257081 produced significant caspase activation [Figure 2 C and E]. However, with the Rho/MRTF pathway inhibitor alone, caspase activation was transient – peaking at about 24 hours. The combination of Vem and CCG-257081 produced strong and <u>sustained</u> apoptosis as detected by both caspase activation and PI staining [Figure 2 C and E]. Similar time courses were obtained when the data were normalized for cell density [supplementary Figure S1]. The effect of the combination treatment on the amount of PI staining in YUMMER_P cells was significantly greater than that of either Vem or CCG-257081 alone. This supports a synergistic mechanism where the two together give a larger effect than the sum of the two individual effects and the sustained apoptosis could result in highly effective cell killing that could limit the development of resistance.

**Figure 2.**
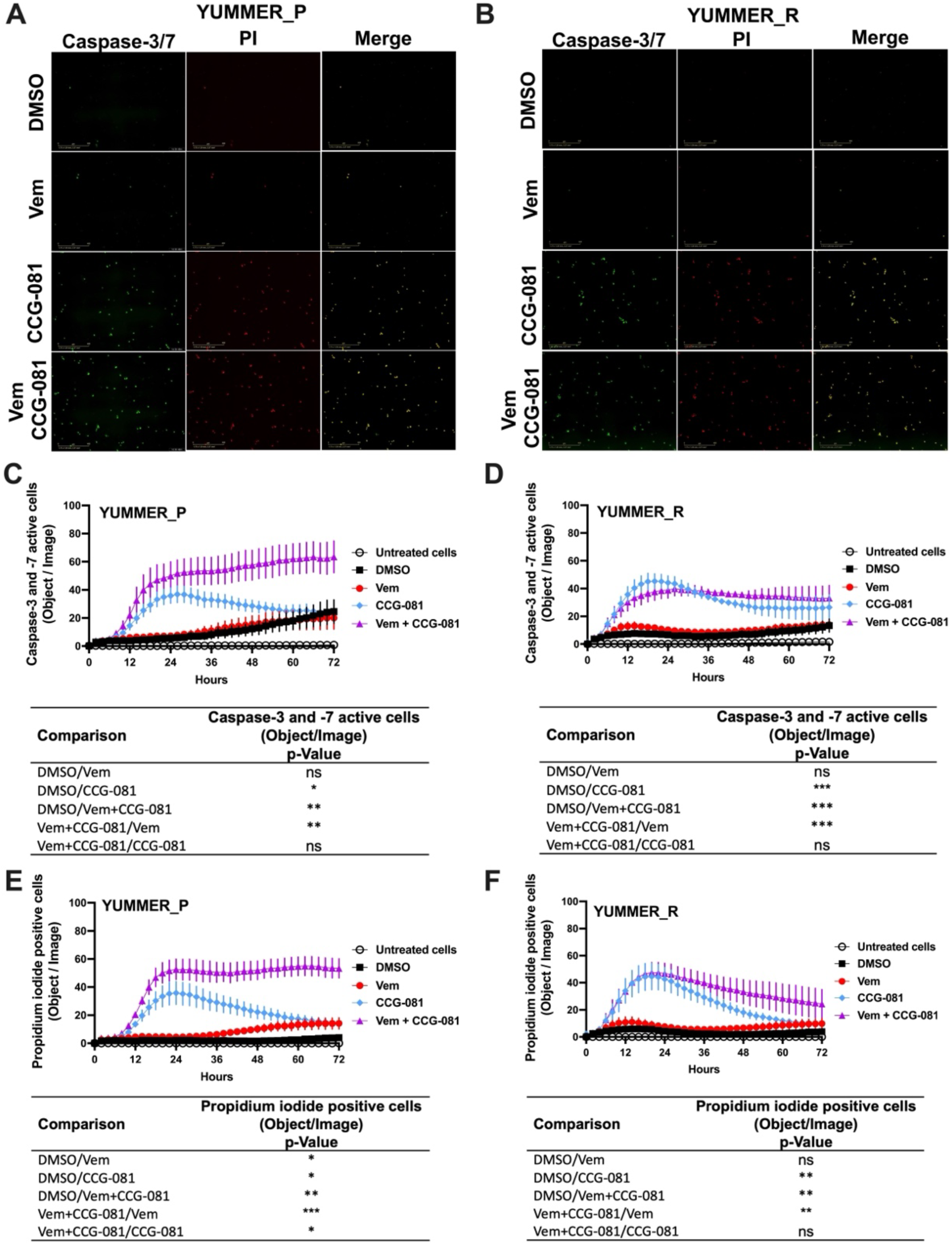
Combined treatment with Vem and CCG-257081 produces sustained apoptosis in YUMMER_P and YUMMER_R cells. Cells were plated and treated as in Figure 1. Treated cells were concurrently incubated with IncuCyte caspase-3/7 Apoptosis Assay Reagent (Green) and Propidium Iodide (Red) as described in Methods. Untreated cells were used as a control. Cells were monitored in real time using the Incucyte S3 live-cell imaging system. (A & B) Incucyte S3 live-cell fluorescence imaging at 24 h. (C & D) caspase-3/7 green readout over time. (E & F) Propidium iodide readout over time. Results are the mean ± S.E. of four independent experiments, * p < 0.05; ** p < 0.01; **** p < 0.0001 by A two-way ANOVA test; ns: not significant.

For the YUMMER_R cells that had previously been made resistant, the Rho/MRTF pathway inhibitor, CCG-257081, did induce apoptosis [Figure 2 D and F], but there was a minimal effect of Vem alone, and, as might be expected, the synergism of Vem with CCG-257081 on induction of apoptosis was lost. Also, the induction of apoptosis was less robust, and the PI detection readout was transient [Figure 2 D and F].

### Inhibition of the Rho/MRTF pathway activates cleavage of the PARP mechanism

Poly (ADP-ribose) polymerase (PARP) is a DNA repair enzyme activated under aberrant cellular changes ^[29–31]^. Cells activate PARP to survive and resolve DNA lesions induced by factors such as anticancer agents^[32–34]^. In apoptotic cells, caspase-3 and -7 have been reported to inactivate PARP through cleavage, inhibiting PARP’s DNA-repairing activities^[35–37]^. To further document apoptosis in our system, we investigated cleaved PARP in YUMMER cells treated for 24 hours with Vem, CCG-257081, or co-treated with Vem and CCG-257081. In the parental YUMMER_P cells, cleaved PARP increased with all three drug treatments, confirming apoptosis induction [Figure 3A]. As expected from the caspase and PI signals, cleaved PARP was not detected in YUMMER_R cells treated with Vem alone. This further confirms the Vem-resistance of these cells [Figure 3B]. Interestingly, cleaved PARP was clearly detected in YUMMER_R cells treated with CCG-257081 alone, but minimal to no cleaved PARP was detected after Vem/CCG-257081 cotreatment. The reason for this is unclear, but perhaps cells with activated PARP died with the combined treatment and weren’t detected. These results show that inhibition of the Rho/MRTF pathway by CCG-257081 activated the caspase-3/7: PARP mechanism in YUMMER cells and remained active in the Vem-resistant YUMMER_R cells.

**Figure 3.**
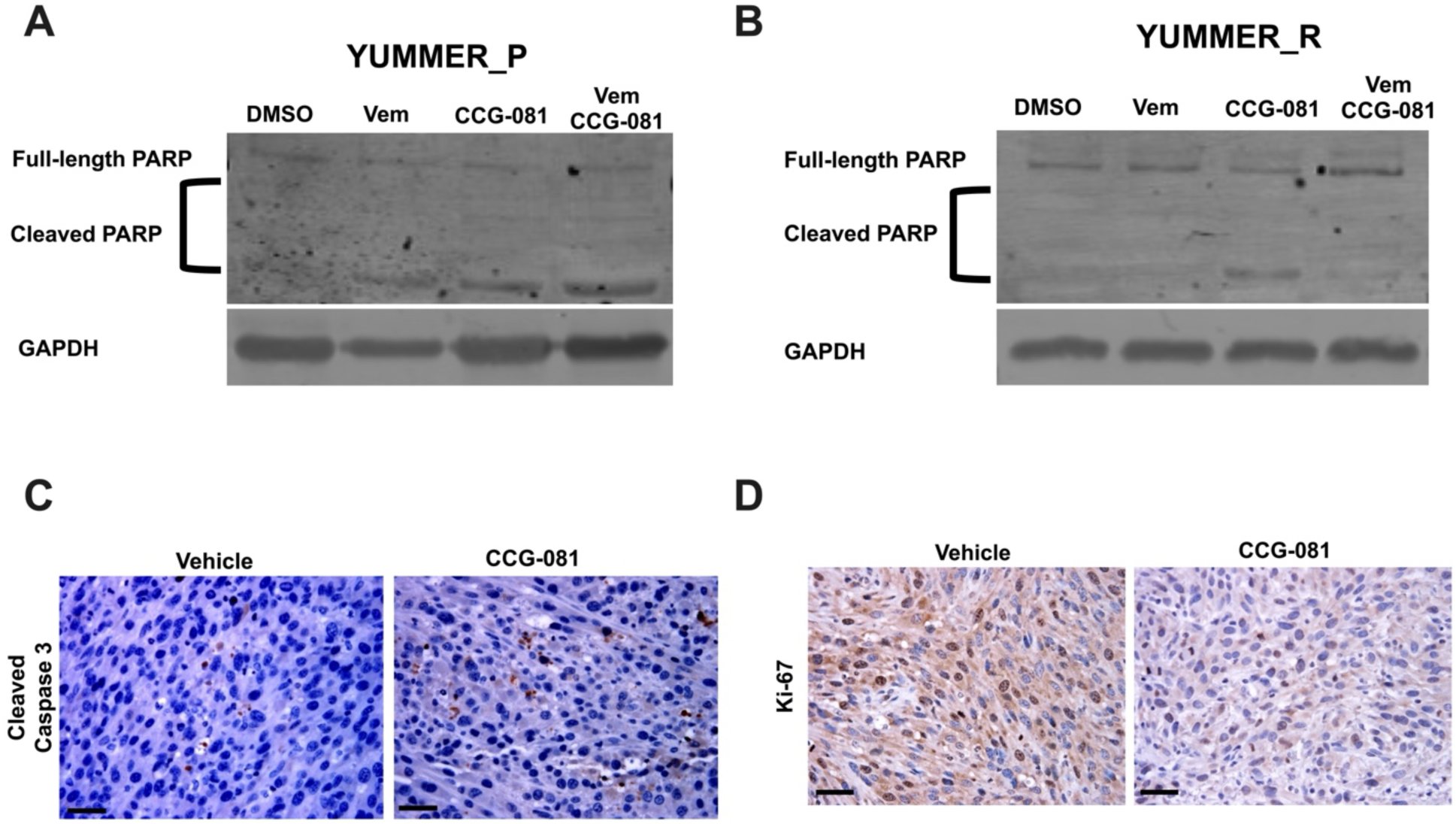
Inhibition of the Rho/MRTF pathway induces apoptosis in YUMMER cells *in vitro* and in YUMMER_R tumors *in vivo*. A-B: Immunoblotting analysis of full-length PARP and cleaved PARP in YUMMER_P (A) and YUMMER_R cells (B) Cell lysate with 30 µg of total protein was loaded, and GAPDH served as loading control. (C-D) Immunohistochemistry staining of YUMMER_R tumor tissues collected from mice treated with Vehicle or CCG-257081 (CCG-081) and stained with cleaved caspase3 antibody (C) or Ki67 (D); scale bar is 5 mm. The images represent data of tissues collected from three tumor-bearing mice injected with the vehicle and three mice treated with CCG-257081.

### CCG-257081 blocks proliferation and induces caspase-3 activation in Vem-resistant melanoma tumors *in vivo*

The strong effect of the pirin-binding, MRTF/SRF pathway inhibitor, CCG-257081, in inducing apoptosis *in vitro* led us to ask if it is also applied *in vivo*. We used the Vem-resistant YUMMER_R cells to develop melanoma tumors in isogenic wild-type C57Bl/6 mice^[21]^. Tumor-bearing mice were treated with a vehicle or CCG-257081 for 12 days, then we collected melanoma tissues. Samples from our previously published study^[23]^ were stained for the proliferation marker a cleaved caspase-3 [Figure 3C] and Ki-67 and [Figure 3D]. Ki-67 staining was reduced in tissues from CCG-257081-treated mice compared to those collected from vehicle-treated mice, showing that the compound reduced cell proliferation in the tumors. CCG-257081-treated tissues also showed higher cleaved caspase-3 staining than the tissues collected from vehicle-treated mice. These results indicate that inhibition of the Rho/MRTF pathway by CCG-257081 induced apoptosis *in vivo* and was associated with reduced tumor cell proliferation.

### Pirin binding and cellular activity of CCG-257081 enantiomers

We recently showed that the pirin-binding MRTF pathway inhibitor, CCG-257081, could prevent development of Vem-resistant melanoma colonies *in vitro*^[12]^. CCG-257081 has two stereoisomers and one publication showed that CCG-1423, the original screening hit from this series, exhibited functional stereospecificity^[38]^. Consequently, we sought to determine whether there was stereospecificity in the binding of CCG-257081 to pirin and in its cellular actions. We tested the two enantiomers (R) CCG-257081 and (S) CCG-257081 for binding to pirin. (S) CCG-257081 had a higher affinity of 8 μM vs >10 μM in a fluorescence polarization competition assay [Table 1]. The IC_50_ for the R enantiomer is probably significantly greater than 10 μM as no inhibition was detected at all at that concentration (the highest tested). The greater affinity of the S enantiomer is consistent with the observation that it was the S enantiomer that was seen in our published co-crystal structure with pirin^[13]^. The S enantiomer also more potently inhibited TGFβ-induced ACTA2 gene expression in human lung fibroblast cells. Surprisingly, there was no difference between the S and R enantiomer, or perhaps even very slightly greater potency of the R enantiomer, in the SRE-luciferase reporter readout of MRTF/SRF-regulated gene transcription in HEK293 cells (Table 1).

**Table 1:**
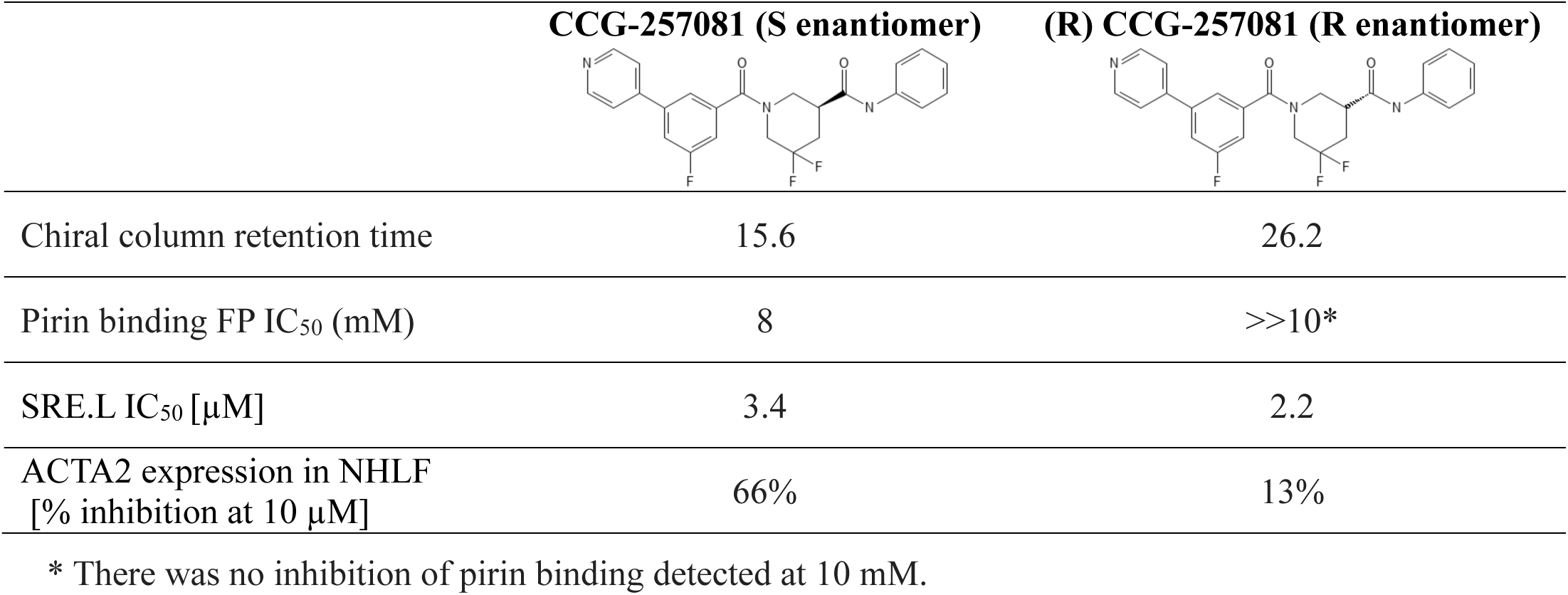
Binding affinities of CCG-257081 enantiomers to Pirin.

|  | CCG-257081 (S enantiomer) | (R) CCG-257081 (R enantiomer) |
| --- | --- | --- |
| Chiral column retention time | 15.6 | 26.2 |
| Pirin binding FP IC<sub>50</sub> (mM) | 8 | >>10* |
| SRE.L IC<sub>50</sub> [μM] | 3.4 | 2.2 |
| ACTA2 expression in NHLF [% inhibition at 10 μM] | 66% | 13% |

\* There was no inhibition of pirin binding detected at 10 mM.

### Prevention of Vem-resistance by CCG-257081 enantiomers

Given the potential for translational use of the pirin-binding-, MRTF-inhibiting compounds for combination therapy with Vem and other MAPK pathway-targeting agents, we asked if there is stereospecificity in the ability of CCG-257081 in preventing resistant colony formation *in vitro*. To mimic a clinical scenario where the compound would be used to prevent the development of resistance to a targeted agent, we cultured the parental YUMMER_P cells for 14 days in the presence of a high concentration of Vem (5 µM). These were the conditions used previously to generate the resistant YUMMER_R cell line^[12]^. Without Vem, the wells would be confluent in only a few days but in the presence of Vem, a modest number of resistant colonies were generated [Figure 4B]. Co-treatment with Vem and different concentrations of the CCG-257081 enantiomers [Figure 4B-D] resulted in significantly fewer resistant colonies. Both enantiomers significantly reduced colony counts at 3 and 10 μM, but only the (S) CCG-257081 significantly reduced colonies at 1 μM. The S enantiomer was significantly more active than the R enantiomer at both 1 and 3 μM. These data indicate that low concentrations of the (S) CCG-257081 enantiomer could potentially be used to inhibit the development of BRAFi-resistant melanoma.

**Figure 4.**
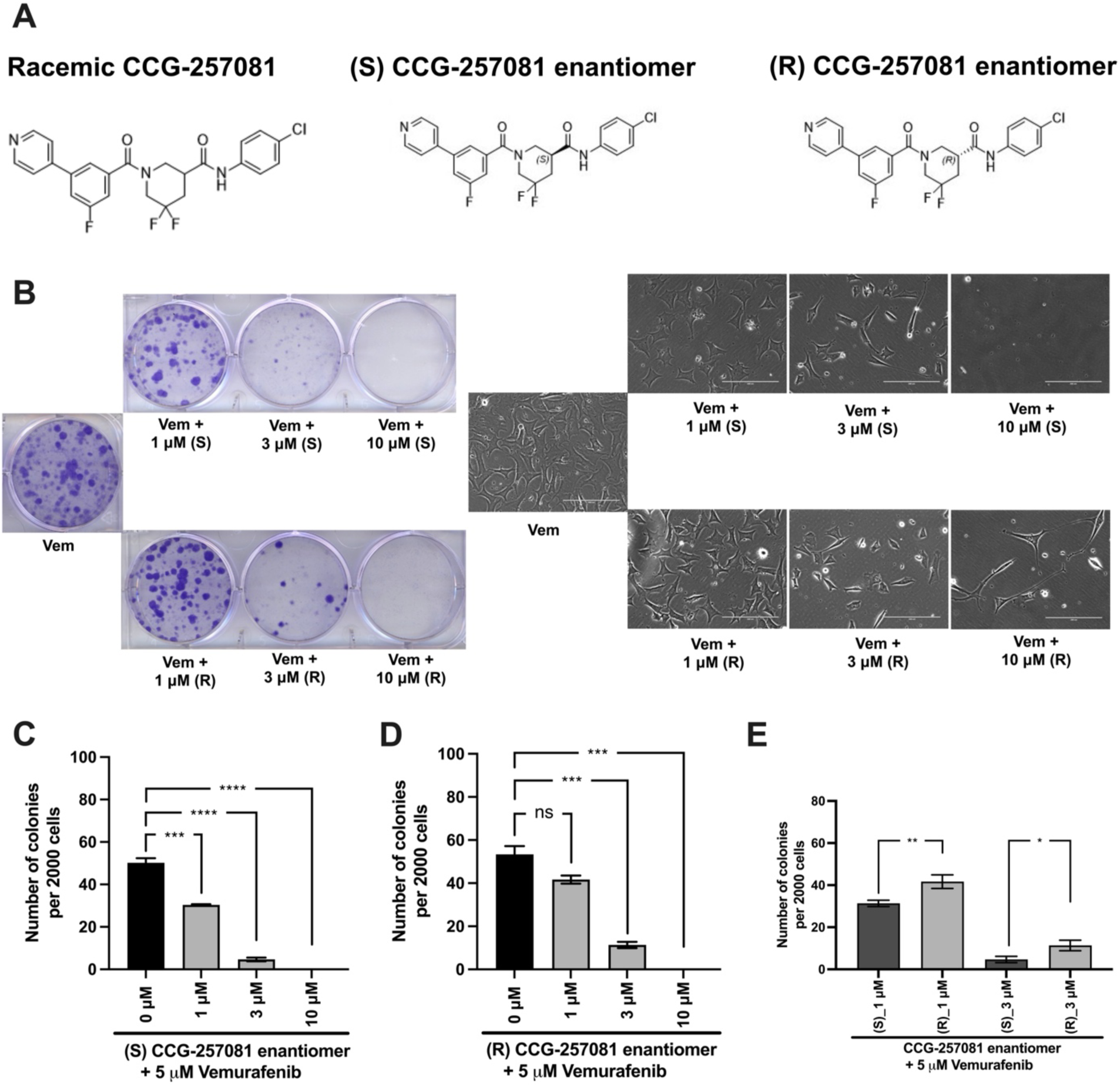
Development of CCG-257081 enantiomers with potent Inhibitory effects on the development of Vem resistance. (A) Structure of the CCG257081 compound and two generated enantiomers, (S) CCG-257081 and (R) CCG-257081. (B) Colony formation assays were done on YUMMER_P cells, cultured in the presence of 5 μM Vem and increasing concentrations of CCG-257081 enantiomers (S and R), as indicated. Colonies were stained with crystal violet, and Images of cells within one colony were captured with a light microscope before staining. C-E: The number of stained colonies was determined using ImageJ. (C-F) Results are the mean of three independent experiments, * p < 0.05; ** p < 0.01; *** p < 0.001; **** p < 0.0001; ns: not significant.

## DISCUSSION

Resistance is a major hurdle facing anti-melanoma therapies – both for targeted drugs and immunotherapy^[1,39]^. Acquired drug resistance develops rapidly in melanoma patients treated with effective targeted therapies such as BRAFi and MEKi. Naïve melanoma cells (not exposed to any treatment) are suggested to include a drug-tolerant cell population.^[40,41]^. These cells are hard to detect because they are scarce and markers are not well-described. However, they expand with drug therapy and develop a stable drug-resistant population. Targeting these pre-resistant cells at the initiation of treatment may enhance anti-melanoma therapies and prevent the development of metastatic melanoma. We previously showed that we could abort the development of Vem resistance in mouse melanoma cells that harbor the prevalent BRAFV600 mutations by treatment^[12]^, *in vitro*, with the pirin-binding Rho/MRTF inhibitor CCG-257081. In about 50% of melanoma cells resistant MAPK-pathway inhibitors, such as Vem, the Rho/MRTF pathway is upregulated upon prolonged drug exposure^[12,28,42–48]^. It is plausible that the pre-resistant cells also upregulate this pathway as a mechanism of resistance. Here, we present a study investigating apoptosis as a potential mechanism whereby CCG-257081 can suppress the onset of Vem resistance.

Previous studies using the Rho/MRTF pathway inhibitor CCG-257081 and related analogs showed promising outcomes in reducing melanoma metastasis^[49]^, enhancing sensitivity to targeted therapies^[49–52]^, and enhancing immune responses to anti-PD1 immune checkpoint therapy^[23]^. Most cancer treatments induce cancer cell apoptosis by activating caspases; such strategy that is central in curing various human cancers^[53–55]^. In the present study, the pirin-binding MRTF pathway inhibitor, CCG-257081, induced activation of caspase-3 and -7 in naïve YUMMER cells (YUMMER_P). Interestingly, Vem alone minimally activated caspase but it synergistically converted the transient activation by CCG-257081 to a sustained activation in a combination treatment of Vem/CCG-257081. Similar results were observed with PI staining and PARP. Thus, we propose that CCG-257081 is targeting a pre-resistant cell population harboring an activated Rho/MRTF pathway while Vem targets the bulk of the cells that have low MRTF activity. The sustained apoptosis likely contributes to the ability of the combination to prevent the development of resistance. For the Vem-resistant cells (YUMMER_R) as well, CCG-257081 induces a transient activation of caspase-3/7 and PI staining however, addition of Vem has a minimal effect. There was, however, a slight, but not significant, trend toward a sustained apoptosis. This is consistent with the reduction of cell growth in the combination treatment but may not be sufficient to completely reverse the resistance.

PARP fixes DNA damage to maintain cell survival, and cancer cells use it to overcome DNA alteration induced by anti-cancer agents^[56]^. PARP inhibition (e.g., inducing PARP cleavage) is a powerful anti-tumor therapeutic tool that stimulates synthetic lethality. PARP inhibitors have demonstrated robust potential in various tumors, such as ovarian cancer^[57]^, breast cancer^[58]^, and advanced melanoma^[59]^. Cleaved PARP has been linked to activated caspases in breast cancer^[36,60]^ and ovarian cancer^[61]^. In our study, we determined a remarkable increase in cleaved PARP that was associated with apoptosis after CCG-257081 treatment. Vem-treated YUMMER_P showed PARP cleavage and insignificant changes in caspase-3/7, referring to other caspases that would regulate cell apoptosis.

Since CCG-257081 has two enantiomers, it is important to know which is mediating the effects on melanoma. Also, this may be a way to help understand the role of pirin in the anticancer actions of these compounds. While our compound clearly binds to pirin^[13]^, the connection between pirin binding, MRTF pathway inhibition, and cancer cell inhibition remains unclear. The S enantiomer of CCG-257081 binds better to pirin in a biochemical assay (Table 1) and that same enantiomer was the one found in a crystal structure of pirin incubated with racemic CCG-257081^[13]^. The S enantiomer is also more potent in blocking TGFβ-mediated activation of ACTA2 gene expression in fibroblasts. ACTA2 transcription is a direct target of MRTF/SRF regulation^[62]^, and its activation by TGFβ is mediated by Rho^[63]^. Interestingly, pirin has recently been shown to contribute to anti-apoptotic effects in colorectal cancer^[13–15,17–20,64]^. While pirin knockdown reduces and pirin overexpression enhances MRTF-regulated gene transcription^[13]^ it is unclear if pirin’s biological effects are mediated by effects on Rho/MRTF or through other mechanisms. Similarly, we cannot be sure whether the effect of the CCG compounds on MRTF signaling is entirely mediated through pirin binding.

Both enantiomers of CCG-257081 showed robust effects on the formation of Vem-resistant colonies. The (S) CGG-257081 was modestly more potent than the (R) CGG-257081 in preventing the development of resistance. This is consistent with the modest difference in activity of the S enantiomer in binding and in suppression of TGFβ-induced ACTA2 expression. While this does not prove that the CCG-compound’s anti-resistance action is through pirin, it is at least consistent with that conclusion. Pirin is involved in various cellular activities and has been implicated in tumorigenesis and malignancy of various cancers^[18,19,64,65]^. Additionally, pirin controls melanoma cell proliferation by regulating the mitochondrial slow-cycling JARID1B gene^[17]^. Slow cycling is characteristic of aggressive melanoma cells, which have the tendency to metastasize^[66]^. So, it is plausible that the compounds are indeed acting through pirin.

Beyond the *in vitro* studies, our recently reported *in vivo* measures of compound effects on the resistant YUMMER tumors^[23]^ provide a preclinical measure of their potential in melanoma therapies. Here, we extend those findings with the measurements of proliferation (Ki67) and apoptosis (caspase 3 activation) showing that apoptosis may well contribute to the ability of the compound to slow the growth of the resistant melanoma *in vivo*. It will be very interesting to know if pirin binding also contributes to the immune therapy enhancement seen with the compound.

The results of the present study propose a model of how CCG-257081 would induce apoptosis of melanoma cells [Figure 5]. Naïve melanoma cells are proposed to include a small fraction of pre-resistant cells that are tolerant to Vem. Upon exposure to Vem and CCG-257081, sensitive cells are killed by Vem while pre-resistant cells are targeted by CCG-257081; we term this Phase I. Cells that escaped from drug selection may differentiate into more drug-insensitive cells (phase II). In phase II, the differentiation might be induced by either Vem or CCG-257081, which is supported by the prolonged activation of caspase-3/7 and extended apoptosis in the co-treated naïve cells. These extended apoptotic events were not observed where the naïve cells were treated with only CCG-257081 or in stably resistant cells cotreated with Vem and CCG-257081. This model suggests that CCG-257081 targets drug-tolerant cells and prevents them from surviving with longer exposure to a BRAFi and inhibits their conversion into stable-resistant cells.

**Figure 5.**
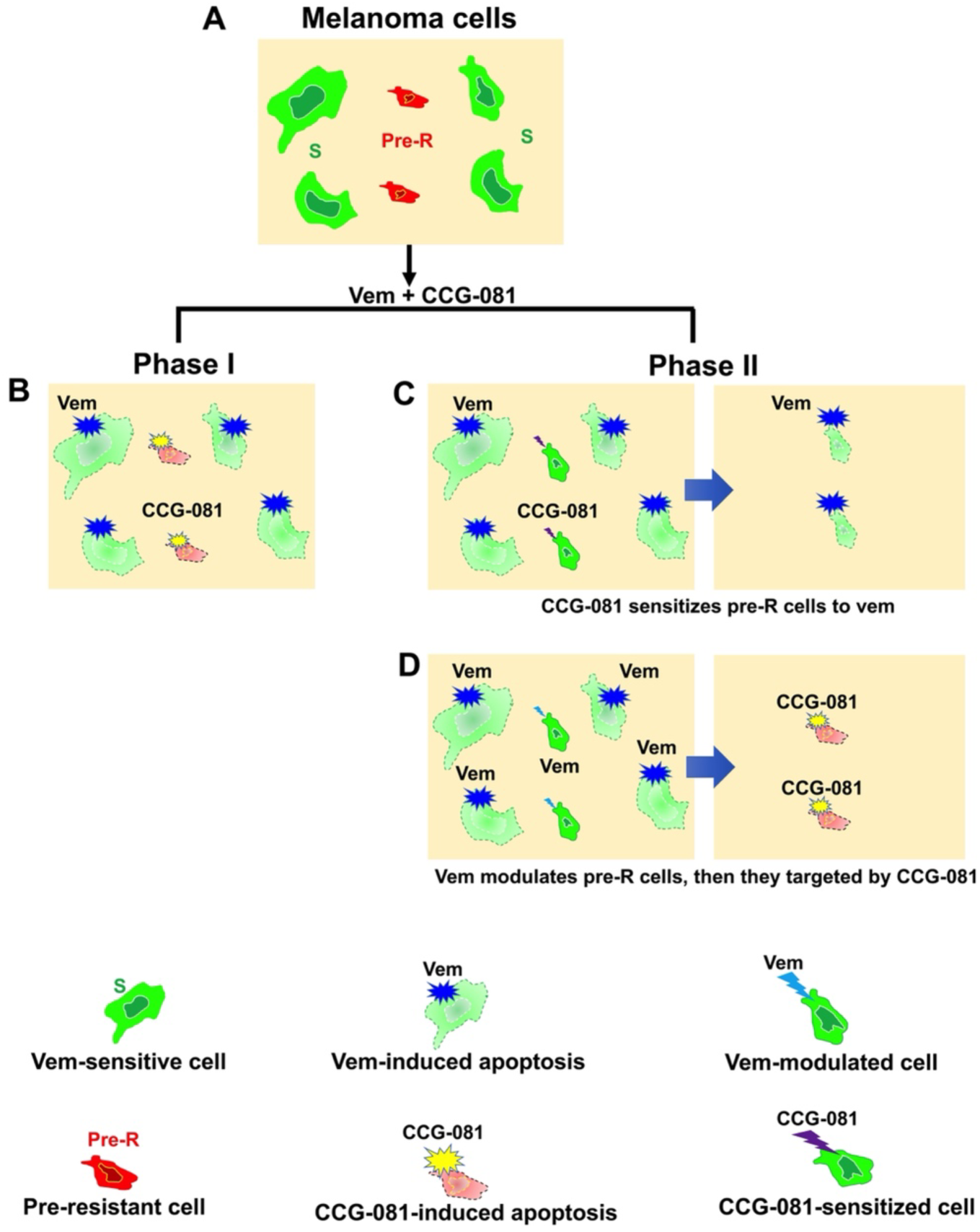
Proposed model of CCG-257081 mediated apoptosis and prevention of drug resistance. (A) Melanoma cells are a heterogeneous pool of predominant Vem-sensitive (S) and rare Vem-pre-resistant (Pre-R). (B) Phase I includes Vem-sensitive cells targeted by Vem. Pre-R cells would exhibit an activated Rho/MRTF pathway that makes them susceptible to CCG-257081, which induces apoptotic events. Two potential models could explain subsequent effects: (C) Phase II results in modulation of Pre-R cells by CCG-257081, converting them into Vem-sensitive cells or (D) Phase II may include cells that are modulated by Vem toward being terminally resistant cells, resulting in cells with promoted Rho/MRTF activities and would then be targeted by CCG-257081 before differentiating into stable-resistant cells.

## DECLARATIONS

## Acknowledgments

The authors thank Mr. Jeff Leipprandt for assistance with the mouse colony and IACUC protocols and Dr. Stephanie Watts for the use of the Incucyte instrument.

## Authors’ contributions

**Bardees Foda** contributed to Conceptualization, Methodology, Formal analysis, Investigation, Writing—Original Draft, and Writing—Review and Editing. **Annika Baker** contributed to the methodology, Writing—Review, and editing. **Richard Neubig** contributed to Conceptualization, Formal analysis, Writing—Review and Editing, Supervision, and Funding acquisition. All authors have read and agreed to the published version of the manuscript. The work reported in the paper has been performed by the authors unless clearly specified in the text.

## Availability of data and materials

Data supporting our study’s findings are available from the corresponding author upon request.

## Ethical approval and consent to participate

Animal studies were performed in accordance with protocols approved by the Michigan State University Institutional Animal Care and Use Committee (IACUC).

## Consent for publication

Not applicable.

