## Supplemental Figure 1 for "Mechanistic insights into Rho/MRTF inhibition-induced apoptotic events and prevention of drug resistance in melanoma: Implications for the involvement of pirin"

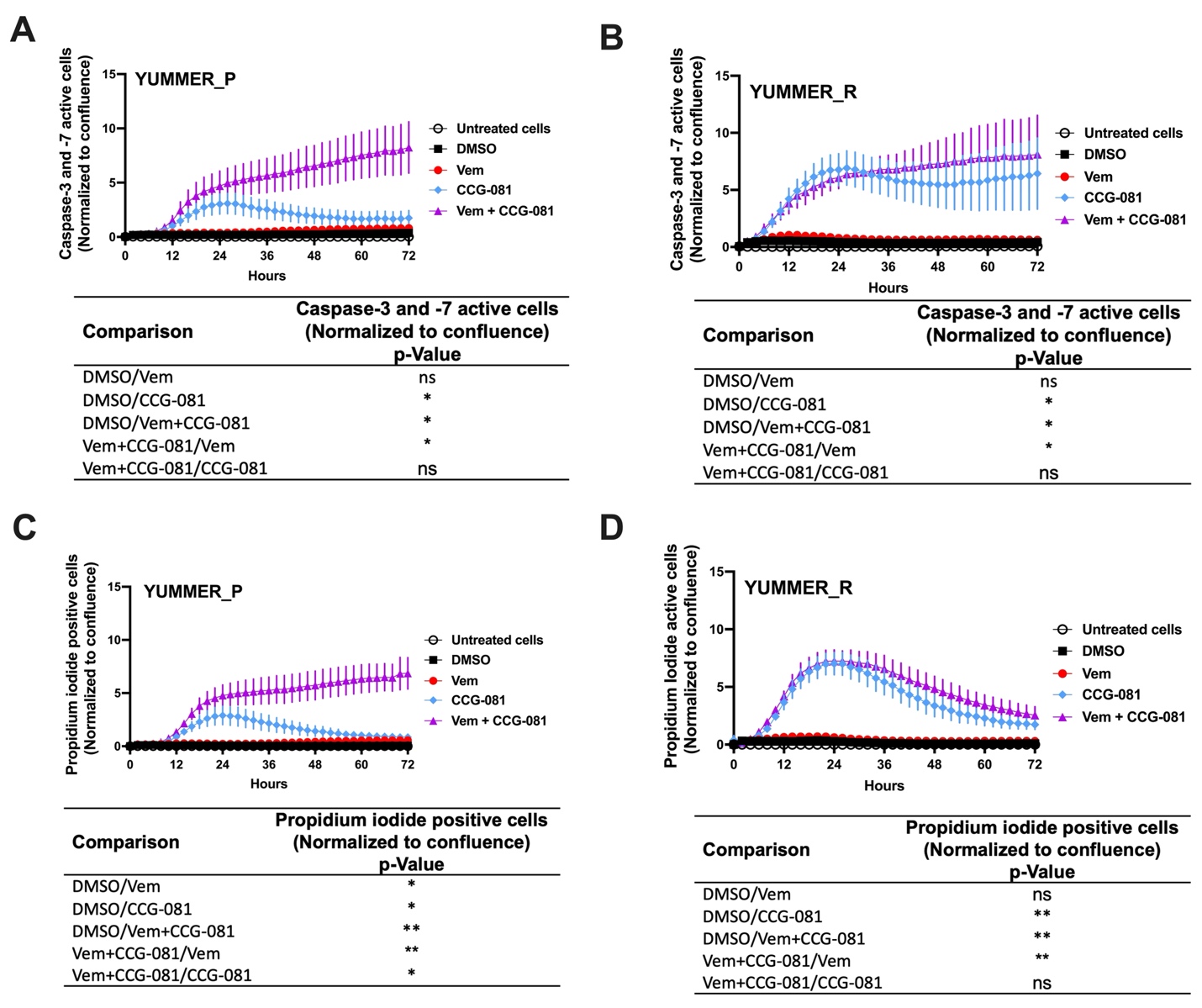
**Supplementary Figure S1.** Activated caspase-3/7 and PI readouts normalized to confluence confirm sustained apoptosis in YUMMER_P and YUMMER_R cells subjected to combined treatment of Vem and CCG-257081 produces. Vem-sensitive melanoma cells, YUMMER_P, and Vem-resistant cells, YUMMER_R, were plated at 1000 cells per well in 96-well plates, and growth was monitored in real-time using the Incucyte S3 live-cell imaging system. The indicated compounds (DMSO control, 10 µM CCG-257081 labeled CCG-081, 5 µM Vem, or both 5 µM Vem and 10 µM CCG-081) were added, and the confluence of the YUMMER cells was calculated using the IncuCyte phase object module. The caspase-3/7 and PI results were normalized to cell confluence. (A & B) Caspase-3/7 green readout over time normalized to cell confluence. (C & D) Propidium iodide readout over time normalized to cell confluence. Results are the mean ± S.E. of four independent experiments, * p < 0.05; ** p < 0.01; **** p < 0.0001 by A two-way ANOVA test; ns: not significant.
